# Alzheimer’s Disease Classification Accuracy is Improved by MRI Harmonization based on Attention-Guided Generative Adversarial Networks

**DOI:** 10.1101/2021.07.26.453862

**Authors:** Surabhi Sinha, Sophia I. Thomopoulos, Pradeep Lam, Alexandra Muir, Paul M. Thompson

## Abstract

Alzheimer’s disease (AD) accounts for 60% of dementia cases worldwide; patients with the disease typically suffer from irreversible memory loss and progressive decline in multiple cognitive domains. With brain imaging techniques such as magnetic resonance imaging (MRI), microscopic brain changes are detectable even before abnormal memory loss is detected clinically. Patterns of brain atrophy can be measured using MRI, which gives us an opportunity to facilitate AD detection using image classification techniques. Even so, MRI scanning protocols and scanners differ across studies. The resulting differences in image contrast and signal to noise make it important to train and test classification models on multiple datasets, and to handle shifts in image characteristics across protocols (also known as *domain transfer* or *domain adaptation*). Here, we examined whether adversarial domain adaptation can boost the performance of a Convolutional Neural Network (CNN) model designed to classify AD. To test this, we used an Attention-Guided Generative Adversarial Network (GAN) to harmonize images from three publicly available brain MRI datasets - ADNI, AIBL and OASIS - adjusting for scanner-dependent effects. Our AG-GAN optimized a joint objective function that included attention loss, pixel loss, cycle-consistency loss and adversarial loss; the model was trained bidirectionally in an end-to-end fashion. For AD classification, we adapted the popular 2D AlexNet CNN to handle 3D images. Classification based on harmonized MR images significantly outperformed classification based on the three datasets in non-harmonized form, motivating further work on image harmonization using adversarial techniques.

## 1. INTRODUCTION

Alzheimer’s disease (AD) - the most prevalent form of dementia - affects more than 20 million individuals worldwide. The disease is characterized by progressive memory loss, and patients often show impairments across multiple cognitive domains that worsen over a period of many years. Neuroimaging can identify anatomical and molecular brain changes in AD - beyond those typically seen with normal aging - such as accelerated gray and white matter atrophy, build-up of beta-amyloid deposits, and abnormal tau protein aggregates 1. In the U.S. alone, there are over 6 million cases of AD 2 - a number projected to double by the year 2050 3.

In 2018, a new consensus definition of biological Alzheimer’s disease was proposed, based on the presence of abnormal biomarkers such as elevated brain amyloid and tau proteins, and neurodegeneration detected as atrophy on brain MRI. Of the biomarkers considered, the high cost and limited availability of PET, and the invasiveness of lumbar puncture procedures to measure protein in the cerebro-spinal fluid, led to interest in more convenient and less invasive assessments such as brain MRI. A convenient diagnostic test for AD based on conventional brain MRI would be highly beneficial to assist diagnosis and to help screen individuals for drug trials, perhaps as a precursor to more invasive but definitive tests such as PET or direct measures of CSF 4. MRI scans capture the spreading profile of neuronal cell loss progressing from the brain’s medial temporal lobes and hippocampus to the association regions of the cortex, eventually spreading to frontal and primary sensory cortices 5, 6. Various machine learning methods based on convolutional neural networks (CNNs) have demonstrated remarkable performance in a broad range of image classification tasks, from handwritten digit identification to labeling of natural images 7, 8. Even so, CNNs such as AlexNet, VGG and ResNet are trained on vast image datasets, that greatly outnumber the available public datasets of medical imaging data. This calls for techniques that that generalize well across multiple data sources even when trained on smaller datasets (typically hundreds of images).

### 1.1 Relevant Prior Work

Many prior machine learning studies have used brain MRI, often combined with other imaging modalities or genetic data, to distinguish patients with Alzheimer’s disease from age-matched cognitively normal elderly individuals, or from people with other diseases or subtypes of dementia. Unsupervised clustering has also been used to subtype patients with different types of dementia 9, and to predict cognitive decline, such as conversion from mild cognitive impairment to AD over a given time period 10.

Within the class of deep learning models based on CNNs in particular, we recently proposed a multi-task 3D CNN that uses an attention mechanism to classify AD and also estimate a person’s age from their MRI scan, a popular benchmarking task, known as the brain age problem 11.

Variations on this basic CNN architecture include the use of hybrid recurrent neural networks (RNNs) to compile information across series of 2D MRI slices 12 as well as techniques for boosting performance, including data augmentation and pre-training 13.

One very large scale study (using 85,721 MRI scans) 14 applied a state-of-the-art deep CNN, Inception-ResNet-V2, to a sex classification task, and then adapted the pre-trained network to AD classification, achieving over 90% accuracy on 3 public datasets.

Even so, domain transfer is an important issue for such algorithms: MRI scanners and protocols vary, and algorithms trained on data from one scanner can perform poorly when tested on data collected on another scanner, or with a different protocol. Efforts to mitigate this problem include the use of generative adversarial networks (GANs) to adjust images so that they resemble those in a reference set or training set (also known as *adversarial domain adaptation*). GANs can be combined with variational autoencoders (VAEs) to embed the imaging information into a latent space that is designed to be scanner-invariant; in that case, the adversary (discriminator) in the GAN can be designed to determine what site or scanner the scan came from, and a data embedding can be generated that makes it harder for the discriminator to determine this, while still retaining features important for other primary tasks, such as classification, and also minimizing a measure of image reconstruction loss. In one such approach, Dinsdale et al. 15 developed a GAN to create scanner-invariant features while simultaneously maintaining performance on a task of interest, which was predicting brain age. Others have used cycleGAN methods on paired data (from a set of people scanned on multiple scanners or with multiple protocols) to interconvert data across scanning protocols 16.

Others have used StyleGANs to separate the content and style of images, and to translate images from one domain to another, without requiring paired training data 17, 18. In a recent innovation, Zuo et al. proposed a multisite image harmonization framework, CALAMITI, that learns a disentangled latent space to interconvert multimodal MRI across 8 scan sites. In their approach, by separating the anatomical representation, which is is scanner-invariant from a contrast representation, which contains scan protocol-specific information, they were able to generate harmonized data for any pair of individuals and scanners.19

In this work, we examined whether adversarial domain adaptation can boost the performance of a CNN model that is designed to classify Alzheimer’s disease. To train and test the method on data with realistic diversity of scanners and populations, we evaluated our method on three public datasets, ADNI, OASIS, and AIBL (detailed in the Methods section). Adopting a recent innovation, we used an attention mechanism to focus the GAN on key features in the image for harmonization; we then translated MRI images from AIBL and OASIS to the reference ADNI format using the Attention-Guided GAN; finally, we compared the performance of our image classification model for classifying Alzheimer’s disease in two sets of data - one consisting of GAN-harmonized images and the other consisting of the original, uncorrected data. Our work is similar in spirit to that of Guan et al. whose *attention-guided deep domain adaptation framework* (AD^2^A) classified patients with Alzheimer’sdisease across 2 public domain datasets using an attention mechanism to identify discriminative regions (as we did in 11) and adversarial learning for domain transfer. Our approach differs in several respects from that paper as we use an attention mechanism to guide domain transfer, rather than to identify discriminative regions for disease classification 20.

## 2. METHOD

### 2.1 Dataset details

In this work, we analyzed three publicly available brain MRI datasets, consisting of standard high quality T1-weighted 3D anatomical scans. The first dataset consisted of scans from the Alzheimer’s Disease Neuroimaging Initiative (ADNI) 21. This included 1,016 subjects, of whom 329 were clinically diagnosed with AD and the remainder were from the control group. The scans include patients primarily from the ADNI2 and ADNI3 studies (second and third phases of ADNI), along with participants from the baseline study (ADNI1) as well as ADNI-GO. The scans were pre-processed by performing reorientation using 22, followed by skull stripping using HDBet 23 which was then followed by bias field correction using *N4BiasCorrection* 24 and then finally performing a 6 degrees of freedom (rigid-body) registration to MNI152 space using FSL 22. The scans were then down-sampled to 2-mm resolution using ANTS 25.

The second dataset consisted of T1-weighted brain MRI scans from the Australian Imaging Biomarker and Lifestyle study (AIBL) 26, which included 628 participants, all clinically diagnosed with AD.

The final dataset was from the Open Access Series of Imaging Studies (OASIS); this dataset included 235 subjects of whom 47 were clinically diagnosed with AD, and the rest were cognitively normal controls.

For harmonization of MRI scans from AIBL to the ADNI format, we selected 620 subjects at random from ADNI and AIBL. In addition, for the conversion from OASIS to ADNI we selected 234 subjects at random from ADNI and OASIS. The above subjects were divided into training and testing sets as per the distribution shown in first part of Table 1.

**Table 1.** Domain adaptation and AD prediction task dataset distributions

|  | **AIBL to ADNI** | **OASIS to ADNI** |
| --- | --- | --- |
| train subjects | 496 | 187 |
| test subjects | 124 | 47 |

|  | **CN** | **AD** |
| --- | --- | --- |
| Training subjects | 700 | 796 |
| Validation subjects | 87 | 99 |
| Test subjects | 88 | 100 |

For the second task, i.e., classification of Alzheimer’s disease, we studied two different scenarios: the first used a combination of scans from ADNI, AIBL and OASIS in non-harmonized form and the other included AIBL and OASIS scans after being converted to ADNI format by the AG-GAN network, along with the original ADNI scans. We used 30 2D slices per scan, selected from specific coordinate positions from the axial, sagittal and coronal sections, as described in section 2.2. The training, testing and validation split was performed randomly and - to ensure a fair comparison - the corresponding training, testing and validation sets for the harmonized and non-harmonized scenarios consisted of identical sets of 2D slices. The dataset distribution by number of patients is shown in the second part of Table 1.

### 2.2 Image Pre-processing

For both harmonization and prediction tasks we used sets of 2D slices from the 3D MRI scans. Based on our prior work comparing fully 3D convolutions (with 3D kernels) to 2D convolutions performed on sets of slices, the latter are more efficient to train and test and lead to more compact models with fewer parameters but comparable performance.

Ten slices were taken respectively from each of the coronal, sagittal and axial sections, yielding 30 slices of 2D images per subject. For the harmonization task, the images were resized to 128*128. However, for our prediction task, we performed multiple experiments by slicing the 3D scans at different indices. As the images were first linearly registered to a standard coordinate space, there is consistent sampling of anatomy across scans, despite some individual variations in anatomy. Hence, ten slices of images around the chosen indexes (5 before and 5 after the index) for each of the three canonical views were selected as sets of 2D images to create our dataset. An assumption of this approach is that the predictive information is not found solely in the discarded slices, which is a fair assumption for Alzheimer’s disease as its effects are diffuse throughout the brain, with some redundancy of information (we test this assumption and evaluate slice selection later in the paper). The images were resized to 224*224 for input to AlexNet. Figures 1, 2 and 3 show examples of the selected 2D slices for participants from ADNI, OASIS and AIBL respectively at specific canonical slice locations. The figures show example slices from the 3 canonical views per scan, as discussed above.

**Figure 1.**
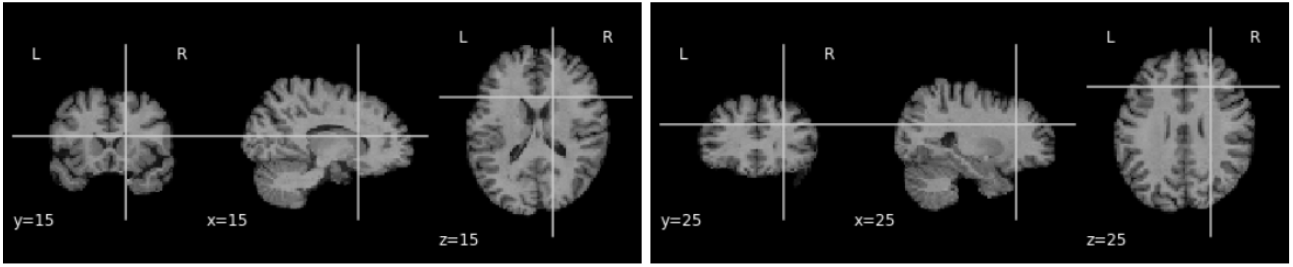
2D slice representation showing y-coronal, x-sagittal, and z-axial views of the brain (MNI template, ADNI scan) when for slices intersecting at (15,15,15) and (25,25,25).

**Figure 2.**
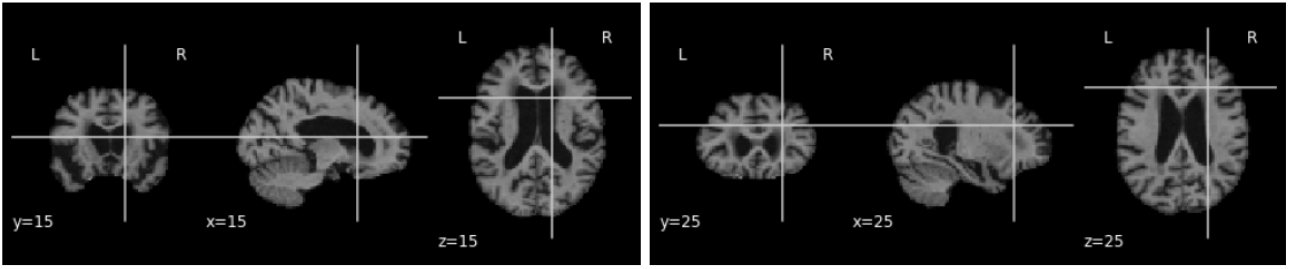
2D slice representation showing y-coronal, x-sagittal, and z-axial views of the brain (MNI template, OASIS scan) when for slices intersecting at (15,15,15) and (25,25,25).

**Figure 3.**
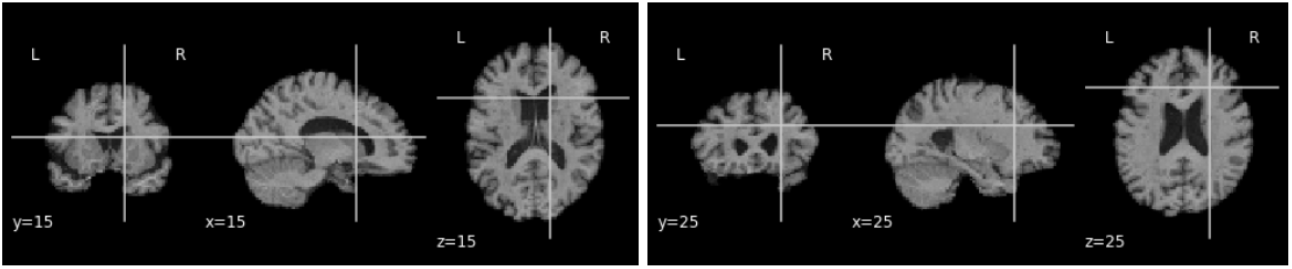
2D slice representation showing y-coronal, x-sagittal, and z-axial views of the brain (MNI template, AIBL scan) when for slices intersecting at (15,15,15) and (25,25,25).

### 2.3 Attention GAN

For our MRI image harmonization task, we adapted the attention GAN model from 17, which was previously introduced to adjust facial expressions (e.g., smiling to not smiling) in 2D color photographs of faces, via style transfer.

Generative adversarial networks are capable of image translation from one domain to another; recent innovations enable this to be performed using unpaired image data. However, the generators may be incapable of identifying the most distinctive parts of the images, leading to low-quality generated images and failures to translate their high-level content. This challenge was handled in 17 using a novel AG-GAN model, also known as an attention *GAN*. A key advantage of this model is the ability to limit undesirable changes to the image content without relying on additional domain-specific information, along with the ability to detect distinctive features across imaging protocols. The AG-GAN uses attention-guided generators which have in-built architecture to generate attention masks and incorporate those with the input image; this, in turn, generates high-quality target domain images.

The authors of 17 also proposed area-specific attention-guided discriminators. The loss functions used by the AG-GAN include attention loss, a pixel loss (defined as an *L1* distance between the input images and the generated images), a cycle-consistency loss and adversarial loss. The model was trained bidirectionally in an end to end fashion. Figure 5 shows the framework outputs given an input domain image.

**Figure 4.**
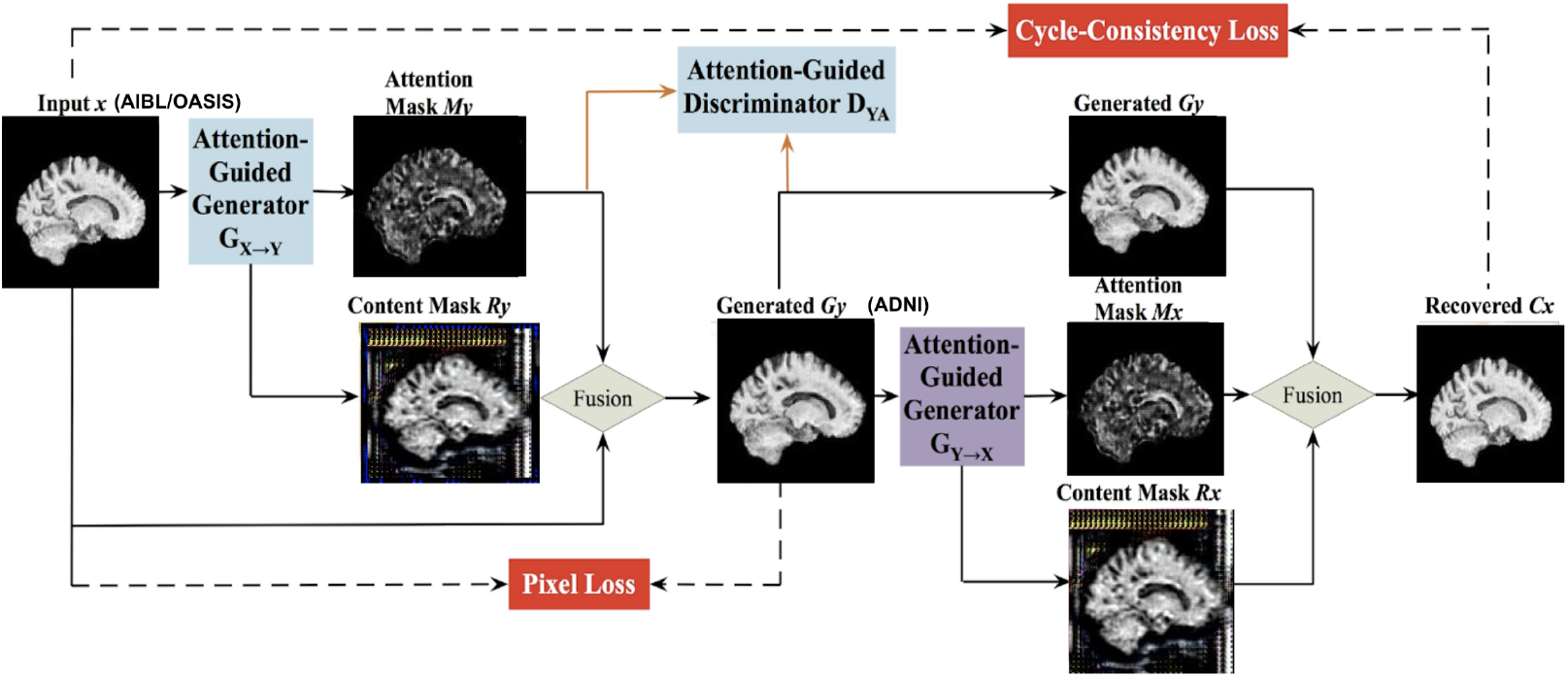
The framework of the Attention-Guided Generative Adversarial Network (AG-GAN) model used here, showing the mathematical transformation mapping for translating subject *x* to subject *y*. There are in-built attention mechanisms in the generators to identify the part of images with maximum distinctions. Architecture adapted from 17, with component panels shown from our neuroimaging application.

**Figure 5.**
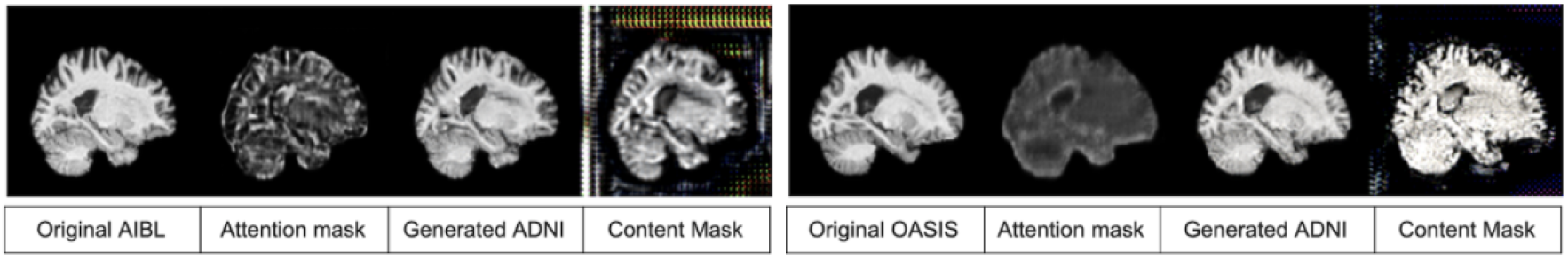
Output of the AG-GAN model given an input domain image along with the attention masks, content masks and the harmonized target domain image

### 2.4 2D AlexNet

AlexNet is a widely-used 2D CNN with fully connected layers. The network contains a total of eight layers; the first five are convolutional and the last three are fully connected. The AlexNet architecture, as shown in Figure 6, is reproduced from 7.

**Figure 6.**
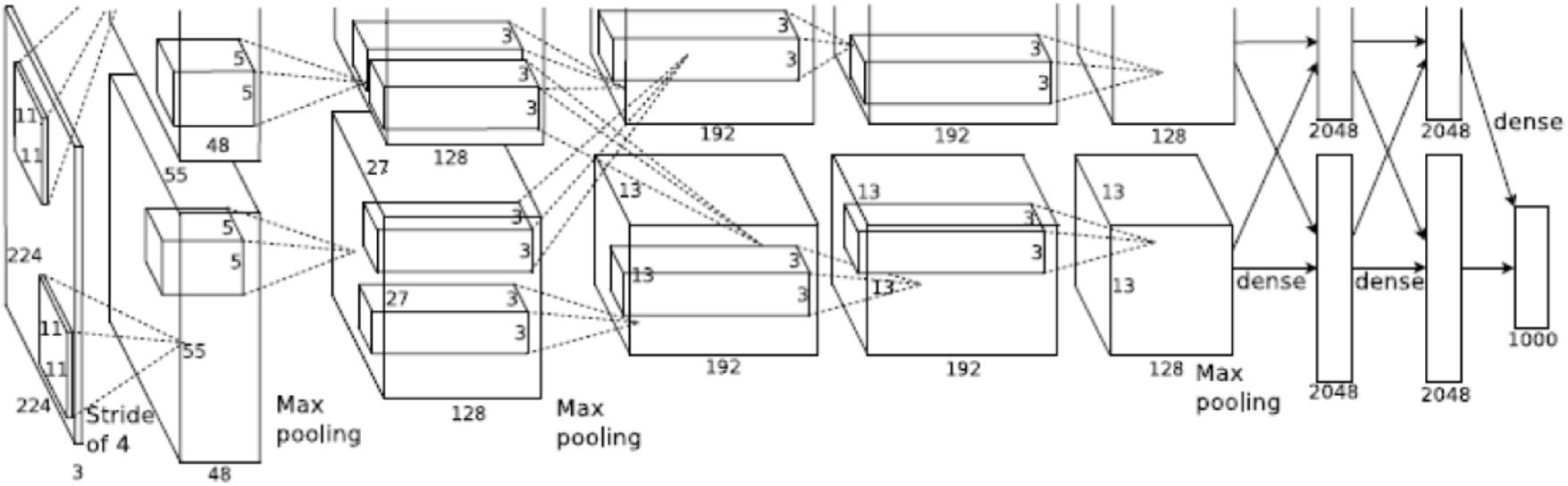
AlexNet architecture

As defined in 7, a 1000-way softmax, capable of reproducing 1000 class label distributions is fed the output from the last fully connected layer. A key factor that makes AlexNet effective is its ability to maximize the multinomial logistic regression objective function. The 3rd convolutional layer’s kernels are connected to all of the kernels from the previous layer whereas to make the network GPU efficient, the 2nd, 4th and 5th layer’s kernels are only connected to those from the previous layer, which were on the same GPU. The fully connected layers had all of their neurons connected to their previous layers. The output from each of the layers have ReLU non-linearity applied to them. The input to the network is an image of size 224*224*3 which is then filtered by 96 kernels with a 4-pixels stride, each having size 11*11*3. As shown in Figure 6, we can see the representation of functionalities among the GPUs. Both of them run separate layer parts at the top and bottom, and interaction occurs only at specific layers.

## 3. RESULTS AND EVALUATION

### 3.1. AIBL to ADNI MRI Harmonization task

The goal of this task is to convert individual images from the AIBL dataset to appear as though they were collected using the ADNI imaging protocol, with comparable contrast and features to the ADNI data, while preserving the anatomical details of the person scanned. We trained the attention GAN network for 100 epochs on the dataset distribution mentioned in section 2.1. Figure 7 and 8 show sample translations for AIBL to ADNI domain adaptation and the training losses respectively.

**Figure 7.**
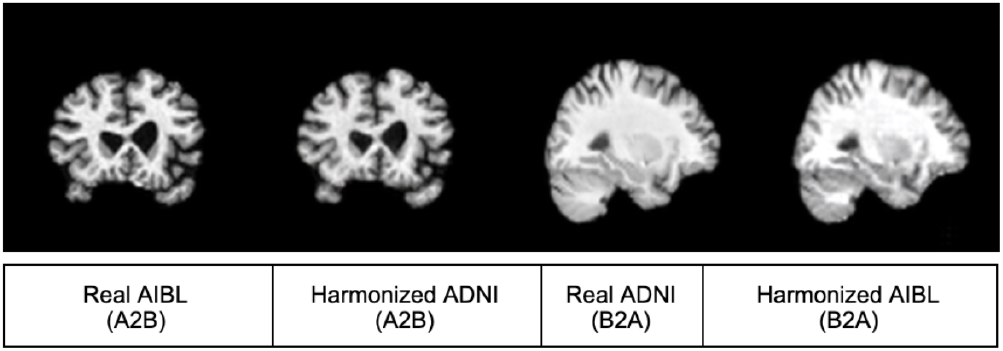
AIBL to ADNI translation sample result

**Figure 8.**
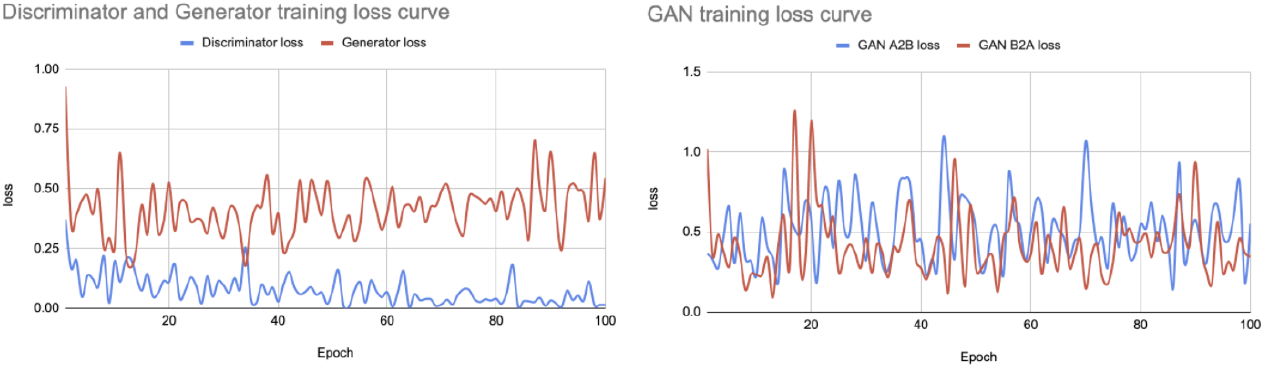
AIBL to ADNI translation loss curve

We evaluated the performance of the GAN models using the metrics listed below.

- SSIM (Structural similarity): compares the similarity between images before and after harmonization to different scans using the structural similarity index measure (SSIM) as a metric. SSIM attempts to model the perceived change in the structural information of the image 27.
- Locality sensitive hashing: uses LSH to find average similarity between images before and after harmonization.

We achieved **0.65** SSIM value for the original AIBL images and **0.66** for the harmonized images which suggests that the structural similarities between the subjects are preserved even after harmonization. Also, the harmonized AIBL images had a LSH value of **0.90** as compared to **0.91** for the original images. The high LSH similarity suggests that AIBL was translated to ADNI format with reasonable style similarity.

### 3.2 OASIS to ADNI MRI Harmonization task

Similar to the AIBL harmonization task, we trained the attention GAN network for 100 epochs on the dataset distribution mentioned in section 2.1. Figure 9 and 10 show sample translations from OASIS to ADNI domain adaptation and the training losses respectively.

**Figure 9.**
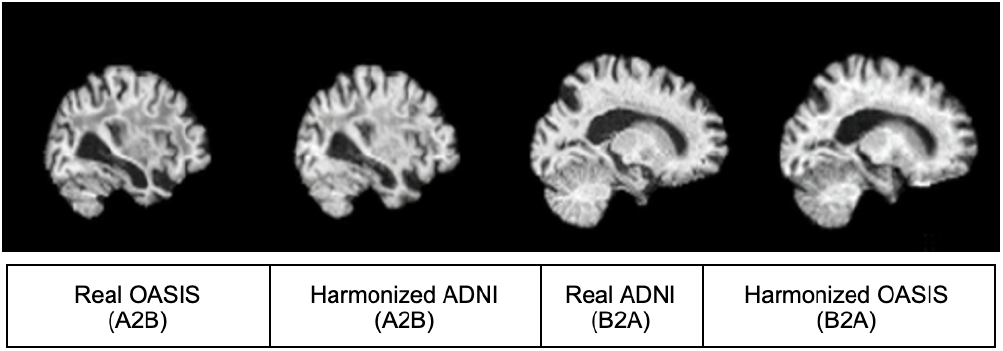
OASIS to ADNI translation sample result

**Figure 10.**
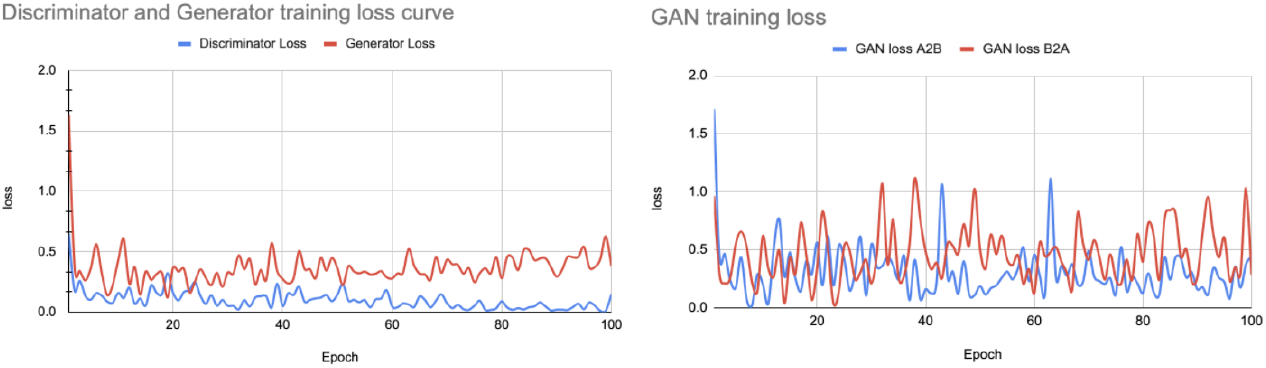
OASIS to ADNI translation loss curve

**Figure 11.**
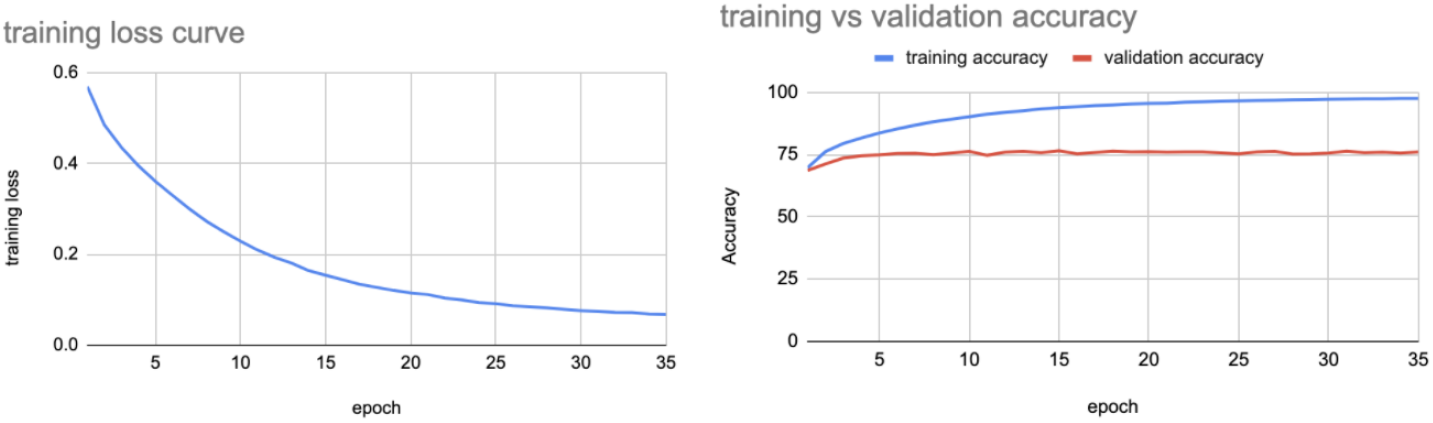
Training loss and training vs validation accuracy curves using non harmonized data for indices 20-59

**Figure 12.**
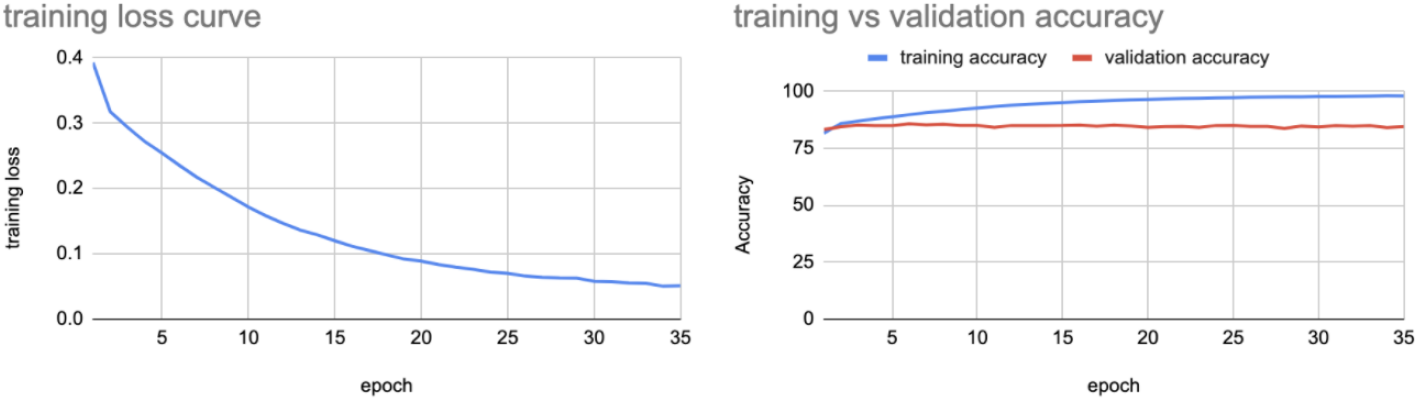
Training loss and training vs validation accuracy curves using harmonized data for slice indices 20-59

We evaluated the performance of the GAN models using the metrics defined in section 2.5.1 and achieved **0.6706** SSIM value for the original OASIS images and **0.6717** for the harmonized images which again suggests that the structural similarities were preserved. Also, the LSH values for harmonized images and original OASIS images were **0.905** and **0.909** respectively thereby ascertaining that OASIS images were translated to ADNI format preserving good style similarity.

### 3.3 Alzheimer’s Disease classification

We trained the AlexNet model defined in section 2.4 for classifying Alzheimer’s disease from the sets of 2D image slices. We conducted multiple experiments using different sets of input 2D images by slicing the scans at various indices, to evaluate the impact of selecting different slice positions on the performance of the model and to consolidate our hypothesis that domain adaptation improves Alzheimer’s detection using deep learning methods irrespective of the slice index positions. We trained our model for 35 epochs for both harmonized and non-harmonized datasets while maintaining identical hyperparameters for both sets of experiments, to observe the difference in performance as a result of harmonization. Table 2 shows the accuracy of our model in classifying people with Alzheimer’s disease, for both the original and harmonized sets of images.

**Table 2.** Alzheimer’s Disease classification model performance

| Slice index range | Accuracy for original images | Accuracy for harmonized images |
| --- | --- | --- |
| 10:19 | 58.00% | 76.60% |
| 20:29 | 71.74% | 83.20% |
| 30:39 | 73.99% | 84.09% |
| 40:49 | 74.03% | 83.39% |
| 50:59 | 73.82% | 82.87% |
| 20:59 | 76.65% | **84.79%** |

To test whether it improved model performance, we increased the number of 2D image slices from 30 to 120 per scan for training and testing. We chose indexes 20-59 in order to retrieve those 120 slices per scan. From Table 2 we can see that for index range 20-59 the model gives the highest accuracy for both harmonized and non-harmonized data sets.

Figure 9 shows the training loss and training vs validation accuracy for the best performing case, using the non-harmonized set of images for classifying Alzheimer’s disease.

The model when tested on test set images gave an accuracy of **76.65%**. We repeated the same experiment on the corresponding harmonized set of images - consisting of AIBL and OASIS 2D slices translated to ADNI format along with original ADNI images, and we observed the training loss and training vs validation accuracy shown in Figure 10.

The model when tested on harmonized test set images gave an accuracy of **84.79%**. We also performed domain adaptation and performed AD classification on MRI scans registered using 9DOF (allowing scaling rather than rigid-body alignment) with the same dataset distribution as the above experiment and achieved the performance shown in Table 3.

**Table 3.** Alzheimer’s Disease classification model performance after 9DOF registration of scans

| Slice index range | Accuracy for original images | Accuracy for harmonized images |
| --- | --- | --- |
| 20:59 | 71.81% | 84.80% |

The experimental results show that domain adaptation of MRI scans from different sources to a single source increases Alzheimer’s disease classification performance by an average margin of more than **8%** irrespective of the slice index positions, as can be seen from Tables 2 and 3.

We also tested our hypothesis for both the brain hemispheres and for the range of indices (−70,-30) we achieved the accuracy as shown below in Table 4.

**Table 4.** Alzheimer’s Disease classification model performance using opposite hemisphere for training and testing

| Slice index range | Accuracy for original images | Accuracy for harmonized images |
| --- | --- | --- |
| -70:-30 | 75.60% | 84.20% |

As AIBL comprised of only AD cases, we tested our hypothesis using only ADNI and OASIS, and observed a 2-3% increase in accuracy as shown in Table 5. The marginal increase in accuracy as compared to our previous scenarios may reflect the lower number of OASIS subjects and a significant domain shift with ADNI dominating the dataset.

**Table 5.**
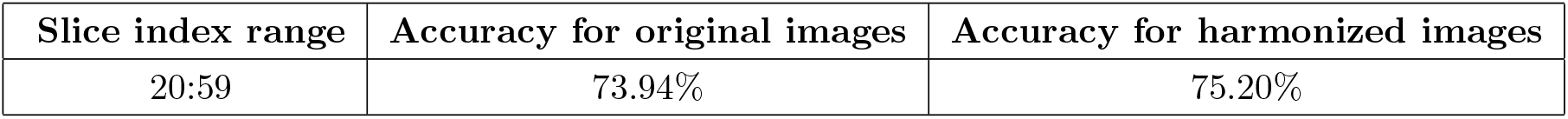
Alzheimer’s Disease classification model performance using only ADNI and OASIS data with ADNI dominating the domain shift

| Slice index range | Accuracy for original images | Accuracy for harmonized images |
| --- | --- | --- |
| 20:59 | 73.94% | 75.20% |

## 4. DISCUSSION

With over 20 million people worldwide suffering from Alzheimer’s disease - a number expected to double by the year 2050, accurate detection of Alzheimer’s disease becomes an utmost priority. Neuroimaging provides us with discernible information on changes in the brain’s macroscopic structure that can be leveraged by image classification models. However, as MRI scanners vary, neural networks trained on data from one source can underperform when tested on data from a different source. It is therefore important to perform domain adaptation across datasets collected from different imaging centers. As scanners differ, as do the scanning protocols employed, the MRI scans generated across sites differ in contrast, resolution, and tissue class differentiation, yielding different intensities that can lead to poor performance in image classification. In this work, we showed that harmonizing MRI scans from different sources to a uniform format significantly improved the classification performance as compared to the case when we use the original, heterogeneous MRI formats. This can help us to merge medical imaging data from multiple sources to learn a better model. This is especially valuable when data relevant to a task is limited. For future work we plan to train our model with even less harmonized data to determine if models based on harmonized data can learn discriminative features more efficiently. We also plan to study how the selection of the reference dataset affects the results. We also plan to compare our method to alternative approaches that perform harmonization directly on 3D MRI scans rather than selected sets of 2D image slices.

## 5. ACKNOWLEDGMENTS

This work was supported in part by the NIH under the AI4AD project grant U01 AG068057.

